# LIDAR explains diversity of plants, fungi, lichens and bryophytes across multiple habitats and large geographic extent

**DOI:** 10.1101/509794

**Authors:** Jesper Erenskjold Moeslund, András Zlinszky, Rasmus Ejrnæs, Ane Kirstine Brunbjerg, Peder Klith Bøcher, Jens-Christian Svenning, Signe Normand

**Author notes:** Corresponding Author: Jesper Erenskjold Moeslund. Contributed equally (Jesper Erenskjold Moeslund, András Zlinszky). E-mail addresses: Jesper Erenskjold Moeslund, András Zlinszky, Rasmus Ejrnæs, Ane Kirstine Brunbjerg, Peder Klith Bøcher, Jens-Christian Svenning, Signe Normand.

## Abstract

Effective planning and nature management require spatially accurate and comprehensive measures of the factors important for biodiversity. Light detection and ranging (LIDAR also known as light radar) can provide exactly this, and is hereby a promising technology to support future nature management and related applications. However, until now studies evaluating the potential of LIDAR for this field have been highly limited in scope. Here, we assess the potential of LIDAR to estimate the local diversity of four species groups in multiple habitat types, from open grasslands and meadows over shrubland to forests and across a large area (approximately 43.000 km^2^), providing a crucial step towards enabling the application of LIDAR in practice, planning and policy-making. We assessed the relationships between the species richness of macrofungi, lichens, bryophytes and plants, respectively, and 25 LIDAR-based measures related to potential abiotic and biotic diversity drivers. We used negative binomial Generalized Linear Modelling to construct 19 different relevant models for each species group, and leave-one-region-out cross validation to select the best models. These best models explained 49, 31, 32 and 28 % of the variation in species richness (R^2^) for macrofungi, lichens, bryophytes and plants respectively. Three LIDAR measures were important and positively related to the richness in three of the four species groups: variation in local heat load, terrain slope and shrub layer height. Four other LIDAR measures were ranked among the three most important for at least one of the species groups: point amplitude entropy, shrub layer density (1.5 – 5 m), medium-tree layer density (10 – 15 m) and variation in biomass. Generally, LIDAR measures exhibited strong associations to the biotic environment, and to some abiotic factors, but was not suitable for representing spatiotemporal continuity. In conclusion, we showed how well LIDAR alone can predict the local biodiversity across habitats. We also showed that several LIDAR measures are highly correlated to important biodiversity drivers, which are notoriously hard to measure in the field. This opens up hitherto unseen possibilities for using LIDAR for cost-effective monitoring and management of local biodiversity across species groups and habitat types even over large areas.

## INTRODUCTION

Nature management typically aims to create, restore, or conserve specific landscape or vegetation structures or natural processes that are favorable for high levels of biodiversity (Polasky et al. 2008, Landis 2017). However, explaining variation in biodiversity across different organism groups and habitats remains a major challenge (Pennisi 2005). While a number of abiotic environmental factors related to soil and hydrology are known to influence local terrestrial biodiversity (e.g., Pharo and Beattie, 1997, Ejrnæs and Bruun, 2000, Moeslund et al., 2013a, Brunbjerg et al., 2017b), the role of biotic resources and spatio-temporal continuity remains hard to quantify and disentangle (Elton 1966, Nordén et al. 2014). Here, we use a comprehensive biodiversity inventory to investigate if airborne Light Detection and Ranging (LIDAR, also termed *light radar*) can adequately represent both the abiotic environment and the biotic factors that shape species-level biodiversity, and hence allow for effective prediction of the variation in local richness of plants, fungi, lichens and bryophytes across a large spatial extent.

LIDAR is increasingly used as a tool for exploring, explaining and predicting biodiversity (Ceballos et al. 2015, Peura et al. 2016, Zellweger et al. 2016, Guo et al. 2017, Vihervaara et al. 2017). Airborne LIDAR records a three-dimensional set of points at sampling densities of 0.1-100 points/m^2^ using a multi-sensor system combining laser ranging, systematic scanning, high accuracy positioning and attitude recording (Wehr and Lohr 1999). Since both terrain- and vegetation surfaces reflect the laser signal, a LIDAR point-cloud includes direct information on both topography and vegetation structure.

The potential of LIDAR-based metrics for investigating and predicting species diversity has already been recognized for local-to-regional scale studies of various species groups (Vehmas et al. 2009, Lopatin et al. 2016, Peura et al. 2016, Thers et al. 2017, Mao et al. 2018). Such studies typically used LIDAR-based indicators of general vegetation structure, such as vegetation height, variance of point height in individual height layers (Froidevaux et al. 2016), and the count (or ratio) of points in various height layers (Vehmas et al. 2009, Mao et al. 2018). Most studies using these vegetation-structure measures were restricted to forests (but see Thers et al., 2017) and have shown that species richness can be modelled with explanatory power up to 66 % for plants and up to 82 % for fungi (Lopatin et al. 2016, Peura et al. 2016, Thers et al. 2017). Instead of vegetation-structure measures, other studies have used terrain measures derived from LIDAR-based Digital Terrain Models (DTM) such as aspect, elevation above sea level, slope, topographic wetness (Moeslund et al. 2013a, Mao et al. 2018) or depth-to-water indices (Bartels et al. 2018). For example, recent work showed that the predictive power of LIDAR-based terrain measures were approx. 20 % and 5–16 % for predicting local plant (Moeslund et al. 2013a) and bryophyte diversity (Bartels et al. 2018), respectively. All the studies that we are aware of, address only one species group or one habitat type and hence none of them are able to generalize their findings across multiple habitat types and species groups. In fact, several of these studies conclude that the next step is to evaluate and validate their modelling results at broader spatial scale and across multiple habitats and species groups (e.g., Peura et al., 2016).

Here, we present a nationwide evaluation of how measures of terrain and vegetation structure – represented by LIDAR measures – can be used to study local biodiversity patterns across multiple terrestrial habitat types and several species groups in Northern Europe. More specifically, we addressed the following questions: (1) To what extent can LIDAR-derived measures (termed “LIDAR measures” hereafter) predict local species richness of vascular plants, macrofungi, bryophytes and lichens across national extent and various habitat types? (2) What are the most important LIDAR measures, and which aspects of the locally measured environment do they represent?

## METHODS

### Study area

Data for this study were collected in Denmark (excluding the island of Bornholm) which has an area of approx. 43.000 km^2^. Denmark is a North European country in the temperate climate zone and is characterized by a lowland landscape (max. elevation ∼170 m above sea level).

### Biodiversity data

Data on biodiversity were collected in the non-winter periods of 2014–2015 as part of a comprehensive biodiversity project covering 130 sites (40 × 40 m) distributed throughout Denmark (Fig. 1) (Brunbjerg et al. 2017a). One hundred of the study sites represented natural and semi-natural habitats. Ten of these were believed to be biodiversity hotspots, while 90 plots were selected by stratified random sampling to cover 5 replicates of 18 combinations of positions along three major natural gradients: fertility (rich, poor), moisture (dry, moist, wet), and successional stage (early, mid, late). Additionally, 15 intensively and extensively cultivated fields as well as 15 managed forest sites were included. For logistic reasons, study sites were clustered into 15 clusters in five regions as showed in Fig. 1. Three sites were left out from analyses either because (1) they were completely inundated during the period where LIDAR data were recorded (causing these data to be erroneous) or (2) their shape was altered by construction works during the biodiversity data collection period.

**Figure 1.**
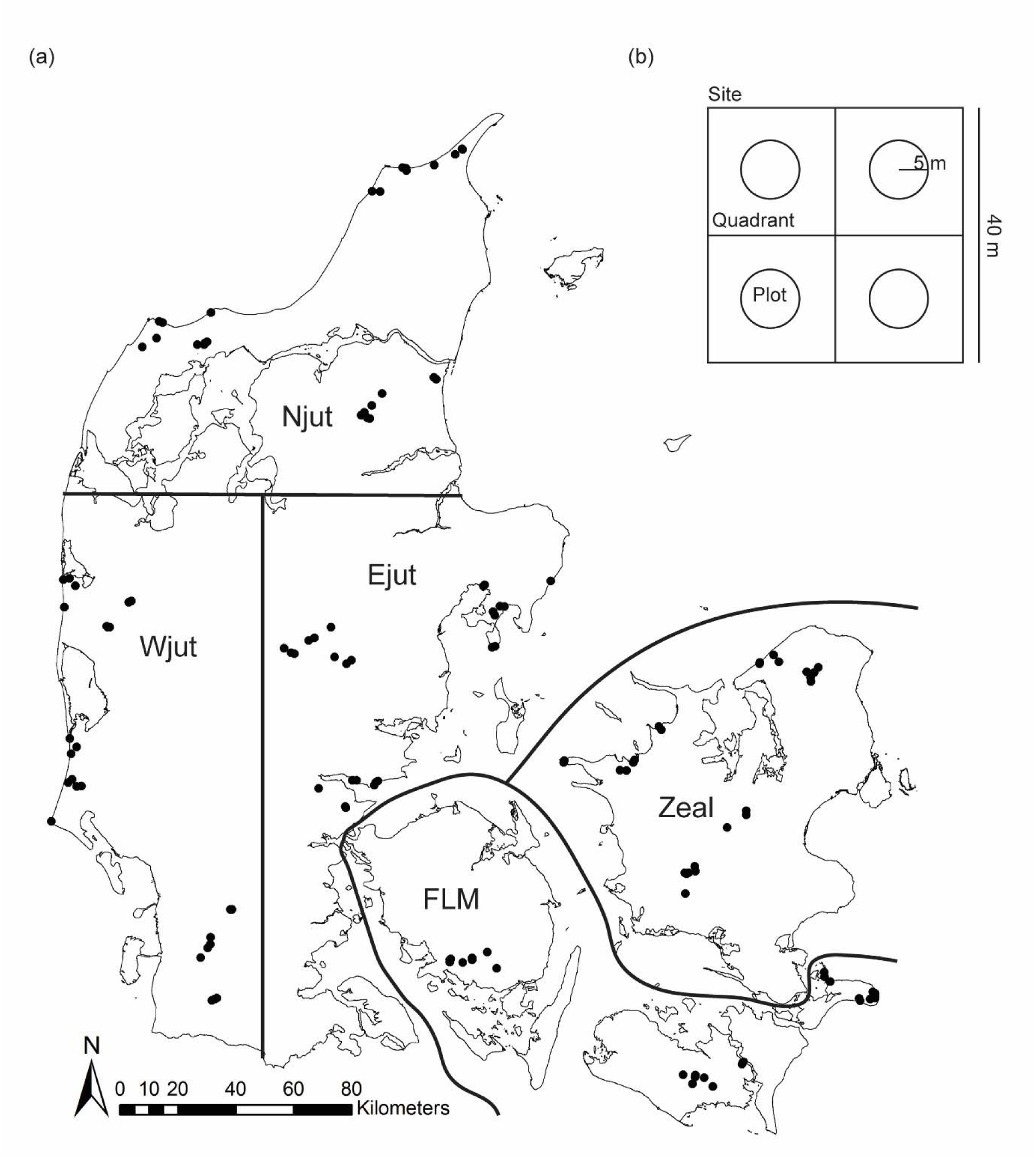
Panel (a) shows a Map of Denmark (excluding Bornholm) with the location of the 130 study sites grouped into 15 clusters within five regions (Njut: Northern Jutland, Wjut: Western Jutland, Ejut: Eastern Jutland, FLM: Funen and smaller islands, Zeal: Zealand). Panel (b) illustrates the study site layout with four 20 × 20 m quadrants each containing a 5-m radius circular sampling unit. From Brunbjerg et al. (2017a)

Leading experts carefully identified all plant, bryophyte, lichen and macrofungi species found in each of the study sites (Fig. 1). Each site was inventoried once for lichens, twice for plants and bryophytes, and three times for fungi (in the fruiting season, August-November). Each inventory had a duration of up to 1 hour. Subsequently, species not readily identifiable in the field were identified in a lab using appropriate equipment. All details on the collection of biodiversity data can be found in Brunbjerg et al. (2017a).

### LIDAR data

We used the latest nationally covering LIDAR point-cloud dataset collected for Denmark (Danish Ministry of Environment, 2015) to quantify and identify terrain and vegetation structures of importance for local biodiversity. This dataset was recorded using fixed-wing airplanes and Riegl LMS-680i scanners operating in the near-infrared wavelength (1550 nm) in a parallel line scan pattern. The airplanes’ flying height was 680 m above ground level and their speed 240 km/h. The data were collected during the leaf-off season in the spring of 2014 (East Denmark) and the fall, winter and spring of 2014-2015 (West Denmark). The dataset has a nominal minimum point density of 4.6 points/m^2^, except for water areas, and is freely available as 1 × 1 km tiles composed of points from multiple strips (kortforsyningen.dk). In the current study, we also used the gridded (0.4 × 0.4 m) Digital Terrain Model (DTM) and Digital Surface Model (DSM) that were based on the point cloud dataset described above (freely available through kortforsyningen.dk, for details see Danish Ministry of Environment, 2015).

To represent vegetation and terrain structure, we calculated 25 measures based on the LIDAR data described above. With one exception (terrain roughness, Table 1), all measures were computed in 10 m resolution. For each measure (except one: root mean square of echo return number, measure 6 in Table 1), we calculated its average within circles of 20 m radius around the center of each study site. LIDAR data processing was carried out using the OPALS software package v 2.2.0.0 (Pfeifer et al. 2014). The full OPALS script, which also holds the exact settings for each calculation, is available in Appendix S1 in the Supporting Information. All measures and their relevant characteristics are detailed in the following. However, to understand all calculation details please refer to the references given in Table 1, which also provides an overview of the LIDAR measures used in this study.

**Table 1.** Overview of the LIDAR measures considered in this study. The variable number is given for convenience and provides a way to quickly link a measure explained in the main text with the same measure in this table. The “Var” column gives the measure of variance if used in this study. References provide calculation details and more information on each measure.

| No. | Name | Alias | Var | Represents | Hypothesis, biodiversity. |
| --- | --- | --- | --- | --- | --- |
| 1 | *Point amplitude* |  | ENT | Succession, leaf and soil moisture | ...depends on succession and moisture balance |
| 2 | *Number of echoes* |  |  | Leaf Area Index, canopy complexity and number of canopy layers | ...is higher in more complex vegetation communities |
| 3 | *Normalized height* | Normalized Z | RAN | Vegetation height | ...is higher when vegetation height varies |
| 4 | *Tree canopy top height* | Canopy top height |  | Height of tree canopies above 3 m | ...may be higher in taller forests when other factors apply as well. May signify old growth forest which often have high biodiversity |
| 5 | *Vegetation penetrability 1* | Echo ratio | ENT | Vegetation penetrability | ...is lower when vegetation is very dense and higher at intermediate levels |
| 6 | *Vegetation penetrability 2\** | Echo no. RMS |  | Vegetation penetrability | ...is lower when vegetation is very dense and higher at intermediate levels |
| 7–12 | *Layer density* | PCount[height interval] |  | Vegetation density of a given layer | ...is higher in open landscapes and forests with shrub layers |
| 13 | *Local LAI* | Pseudowaveform | VAR | Local leaf area index (LAI) | ...is lower when vegetation is very dense and higher at intermediate levels |
| 14 | *Biomass* |  | ENT | Biomass, litter mass, deadwood | ...may be higher when biomass is high. May also be high in open habitats with low biomass levels |
| 15 | *Crown base height* |  |  | Height of tree crown bases | ...is higher when tree crown base is high as this may indicate old growth forest |
| 16 | *Crown span* |  |  | Vertical extent of tree crowns | ...is higher when crown span is high as this may indicate old growth forest |
| 17 | *Shrub layer height* |  |  | Shrub layer height | ...is higher in forests with shrub layers |
| 18 | *Canopy openness* |  | VAR | Light conditions | ...Is higher when the canopy is not too closed (forests with gaps) |
| 19 | *Terrain slope* | DTM Slope |  | Soil moisture, heat balance, bare soil | ...depends on terrain slope |
| 20 | *Topographic wetness index* | TWI |  | Soil moisture | ...depends on moisture balance |
| 21 | *Heat load index* | DTM Heat | VAR | Heat balance | ...is often lower when the terrain is very dry |
| 22 | *Terrain roughness* | DTM SigmaZ 0.5 |  | Microscale (0.5 m resolution) terrain heterogeneity/roughness | ...may be higher when terrain varies more |
| 23–<br>24 | *Terrain openness* | DTM openness & DTM landscape openness |  | Local and landscape scale terrain heterogeneity | ...may be higher when terrain varies more |
| 25 | *Terrain linearity* | DTM openness difference (min-max) |  | Local terrain pattern linearity | ...is lower when terrain is more linear (human influenced) |

\*: Only the variability measure was calculated and used in this study.
Abbreviations: ENT: Shannon entropy, RAN: Range, VAR: Variance, RMS: Root mean square error, DTM: digital terrain model

#### Vegetation structure measures

To represent succession stage and moisture balance in both vegetation and soil we retrieved the *point amplitude* (measure 1, Table 1) of the points in the LIDAR point cloud. A point’s amplitude will be high if the target reflecting the laser light is flat and has a high reflectivity. It will be low for tall canopies where the light energy is distributed between a number of returns, for complex or opaque surfaces such as leaves, and for surfaces with low reflectivity. At the wavelength used here, vegetation surface reflectivity (and thus point amplitude) relates to leaf water content (Junttila et al. 2018) and soil moisture (Zlinszky et al. 2014).

To reflect canopy complexity and number of canopy layers we retrieved the *number of echoes* (measure 2) returned by each laser pulse emission. Single echoes are returned from continuous surfaces (e.g., flat arable fields) larger than the sensor footprint (the area illuminated by each pulse of laser light from the sensor), while multiple echoes are generated when the pulse hits several surfaces at different distances from the sensor (e.g., a relatively open forest with shrubs, under-forest and trees having leaves or twigs at different elevations). Note that in dense forests, some LIDAR pulses may not penetrate and reach surfaces below the upper parts of the canopy and therefore the number of echoes may be relatively low here. Since the number of echoes correlates with the number of overlapping vegetation layers, it also represents the Leaf Area Index (LAI). In the point cloud dataset used here, the upper limit for number of echoes recorded was five.

To represent vegetation height we subtracted the local DTM from the DSM giving the *normalized height* (measure 3) and subsequently we computed the *tree canopy top height* (measure 4). The latter differs from the first in the calculation procedure. Canopy top height is based on the 90^th^ percentile of points above 3 m and below 50 m (Mücke, Deák, Schroiff, Pfeifer, & Heilmeier, 2014), and is consequently undefined when the surface height is outside this span.

To mirror *vegetation penetrability* (measures 5–6) we calculated the echo ratio and the root mean square error (RMS) of the echo return number. Both measures reflect the penetrability of the canopy (Höfle et al. 2012). Echo ratio is high where the surface is impenetrable and relatively flat and lower where the surface is uneven or penetrable. The RMS of the echo return number is high when the vegetation is relatively tall and dense, and low when vegetation is low or impenetrable.

To reflect vegetation density in different canopy layers we calculated the *layer density* –typically referred to as the point count – in six height intervals above ground (measures 7–12, see also Fig. 2), starting with 1.5–5 meters, upwards in steps of 5 m until 30 m (similar to Zellweger et al., 2014).

**Figure 2.**
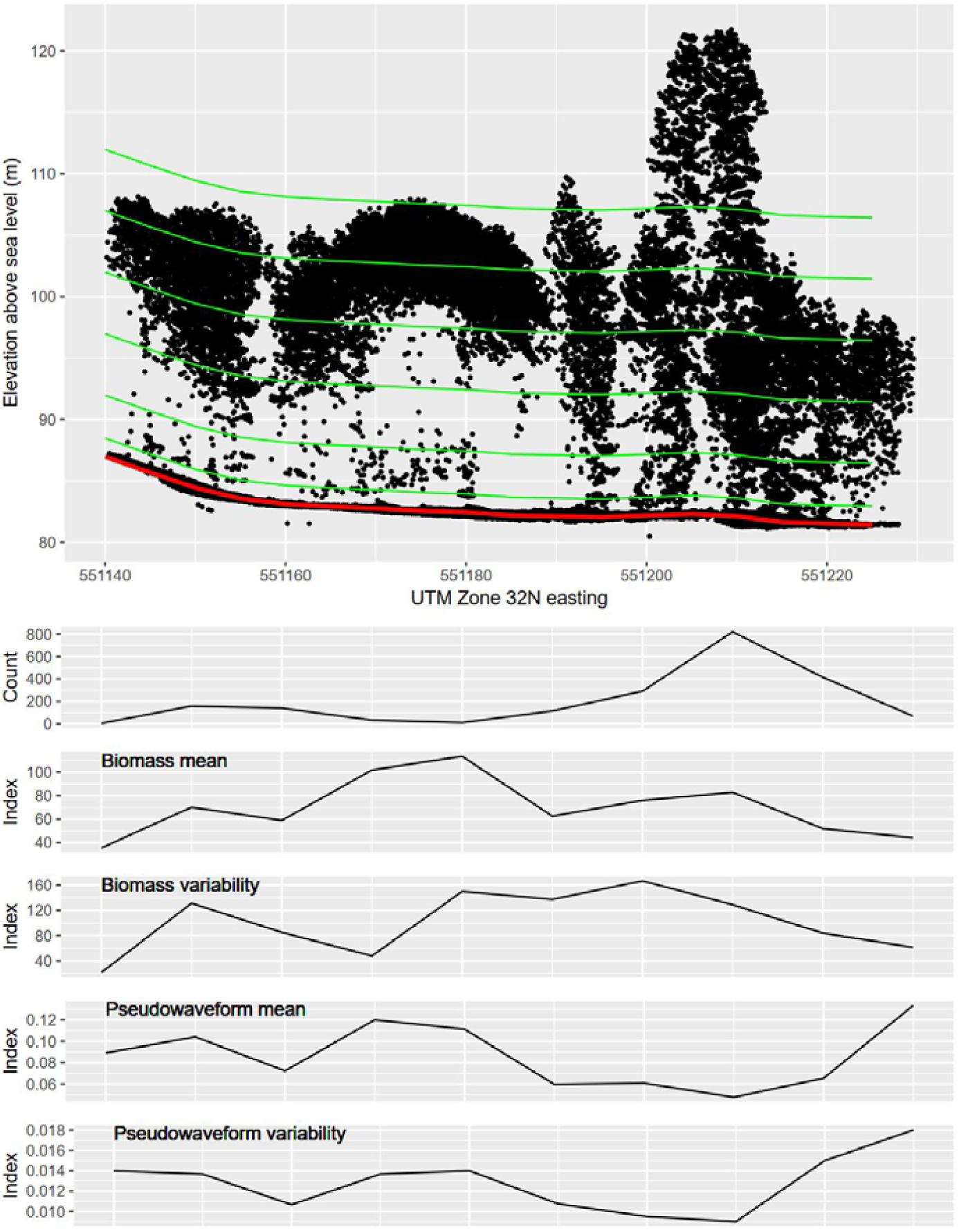
Cross-section of a LIDAR point cloud and examples of LIDAR measures and their variability. The uppermost graph shows the point count, while the remaining four graphs show the values of *biomass* (measure 14) and *local leaf area index* (measure 13) and their variability measures in the cross-section. Each black dot represents a point in the point cloud. Green lines delimit the vegetation layers relative to height above ground used for calculating layer density (measures 7–12). The red line marks ground level.

To approximate the *local leaf area index (LAI)*, we calculated the “pseudowaveform” (measure 13, see also Fig. 2) following Van Aardt et al. (2012). This measure is low when the local LAI is high meaning that the canopy is dense and the LIDAR pulses hardly penetrate the canopy. If LAI is lower, the LIDAR pulse can penetrate further into the canopy giving a higher pseudowaveform-value.

To estimate biomass, we developed a new index of relative *biomass* (measure 14, see also Fig. 2) by calculating a weighted combination of multiple structural attributes based on the recommendations of McElhinny et al. (2005). Biomass correlates with vegetation height, but is also influenced by vegetation density and vegetation layering. Therefore, we combined Normalized height, echo ratio and number of echoes in a weighted sum to create a biomass measure (Eqn. 1). The weighting was selected to obtain a value equal to vegetation height in the simplest cases (large trees, no significant understory), and higher if vegetation is denser or has more layers than in these simple cases.

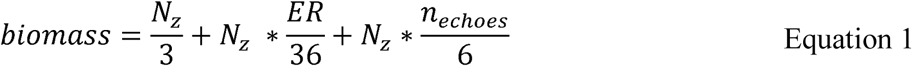

N_z_ is normalized height of the first LIDAR echoes, ER is echo ratio, n_echoes_ is the number of echoes generated by each LIDAR pulse. Before using this measure in our analyses, we checked that it correlated highly with measured factors typically thought to mirror the actual biomass such as litter mass, dead wood volume and vegetation height. This was indeed the case (Table S2)

To represent the height of tree crown bases (i.e. the lowest point of a crown) we calculated the *crown base height* (measure 15). This is based on the 5^th^ percentile of the height distribution of all LIDAR points above 3 m and below 50 m (Mao et al. 2018).

To reflect the sizes of tree crowns we calculated the *crown span* (measure 16). This is the height difference between the canopy top height (measure 4) and the crown base (measure 14).

As an estimation of the *understory height* (measure 17), we calculated the 90^th^ percentile of the normalized heights between 0.3 and 3 m.

To represent light conditions, we calculated *canopy openness* (measure 18) for all points categorized as “ground”, but contrary to terrain openness (see below), we calculated this considering vegetation points as well. Therefore, canopy openness relates to the actual occlusion of sky view of ground points by the canopy around them. Canopy openness is high for ground points inside canopy gaps, and low for ground points beneath a closed canopy.

#### Terrain structure measures

To represent key features of the local terrain (e.g., soil moisture or heat balance, Moeslund et al., 2013b), we calculated *terrain slope* (measure 19) and terrain aspect (used for heat load index calculation, see below) directly from the DTM.

As a proxy for local moisture conditions we used the *topographic wetness index* (TWI, measure 20, Hengl and Reuter, 2009) from Moeslund et al. (2013a). To match the resolution of the rest of the measures we aggregated (average) this TWI layer to 10 × 10 m.

To reflect local heat balance, we calculated the *heat load index* (measure 21, cf. McCune and Keon, 2002) based on terrain aspect. This index reaches maximum values on southwest-facing slopes and zero on northeast-facing slopes.

To estimate local *terrain roughness* (measure 22), we used the points classified in the point cloud as “ground” to calculate sigma Z at a 0.5 × 0.5 m resolution (i.e., a robust indicator of standard deviation) in a search radius of 0.75 m. On purpose, this was the only LIDAR measure not calculated at 10 × 10 m resolution enabling us to test for micro-scale terrain heterogeneity effects.

To represent local and landscape scale terrain heterogeneity, we calculated the *terrain openness* (measures 23–24, also known as sky-view factor, Doneus, 2013) at 10 and 150 m spatial scales (kernel radius). Terrain openness can be calculated for any point of interest. It is defined as the angle of a cone (having the radius of the kernel) turned upside down – with its tip restrained to the point of interest – when it touches the points closest to the surface normal vector. This measure is high in flat (relative to the scale at which it is calculated) areas and low in heterogeneous terrains.

To estimate the *terrain linearity* (measure 25) we calculated the difference between minimum and maximum terrain openness (see above). Maximum openness is high if at least some part of the terrain is open, whereas minimum openness is high when the terrain is open in all directions surrounding the point of interest. In randomly rough surfaces, minimum and maximum openness are quite similar, but in terrain locations with linear features, maximum openness is high (along a ditch or embankment for example) while minimum openness is low (along the sides of a linear terrain feature). Therefore the difference in minimum and maximum openness is high where linear features with a clear direction – typically man-made – occur (Zlinszky et al., 2015).

To enable a test of the importance of variability in the LIDAR measures we calculated a number of variability measures: *standard deviation*, *root mean square error* and *Shannon entropy* and in some cases the *range*. We did this only for LIDAR measures for which we believed it made ecological sense (the measures marked with a variability measure in Table 1).

### Locally measured environmental data

To support the ecological interpretation of our LIDAR measures, we used data for a number of biotic and abiotic factors. These factors were measured or estimated at each of the study sites. The protocols for these measurements and estimates can be found in (Brunbjerg et al. 2017a). We obtained data on the following 15 locally measured or estimated factors for this study: mean difference of day and night temperatures for (1) air and (2) ground surface respectively, (3) median light intensity all year, (4) median soil moisture in May, (5) leaf nitrogen (N), (6) leaf phosphorus (P) and (7) leaf N/P ratio, (8) soil N, (9) soil P and (10) soil pH, (11) litter mass, (12) total basal area of trees larger than 40 cm diameter at breast height, (13) deadwood volume, (14) mean herb layer height and (15) temporal continuity class.

### Data preparation

For a given LIDAR measure, its variability measures (e.g. the root mean square error, Shannon entropy and standard deviation of vegetation height) were always highly correlated (Spearman’s rho > 0.7, Fig. S3). Consequently, for further analysis we retained only the variability measure (for each LIDAR measure) showing the highest mean correlation to the species richness of all four species groups (Table 1 shows which variability measure we retained).

For statistical analysis, we used the species richness of plants, bryophytes, lichens and macrofungi as response variables, modelling each species group individually. We used both the LIDAR measures and their respective variability measures (as showed in Table 1, 25 LIDAR measures and seven variability-measures, in total 32) as predictors in our models. We wished to evaluate the performance of all predictors at least once and consequently divided the predictors into 19 sets of uncorrelated predictors (i.e. Spearman’s rho <= 0.7 with any other predictor in the set) (Fig. S3 and Appendix 4, Tables S4.4–S4.7).

Prior to analysis, the nature of each predictor’s relationship to the response variable was checked visually and the predictor in question was either logarithmically or square root transformed if needed to ensure normality (Table 1). In a few cases (i.e., 2 – 4 depending on species group) this check caused us to suspect quadratic relationships. In these cases, we used Akaike’s Information Criterion (AIC) to evaluate if including the squared term of the predictor improved the model (see *Statistical modelling* below). For this, we used a backward stepwise selection procedure following best practice as recommended by (Burnham and Anderson 2002). Since this evaluation did not reveal any quadratic relationships, we continued modelling using only linear terms.

### Statistical modelling

#### Explanatory power of LIDAR-based measures for local species richness

We used Generalized Linear Models (GLMs) to examine the explanatory power of the high-resolution airborne LIDAR-based measures for local species richness. Species richness (count data) is usually expected to follow a Poisson distribution. However, initial implementation of GLMs with a Poisson error distribution and logarithmic link function were overdispersed. Therefore, we used negative binomial GLMs.

To select the overall best model between the set of 19 candidate models for each species group, we conducted a fivefold leave-one-region-out cross validation for every candidate model. Thus, for each region (see *Biodiversity data* and Fig. 1) we predicted species richness using models calibrated on data from the other four regions. We used non-parametric rank correlation (Spearman’s rho) between predicted and observed values to select the best model for each species group. This procedure was adopted to secure robust model selection with respect to overfitting, potential multi-collinearity and spatial autocorrelation. During model selection, we did not encounter issues with non-normally distributed model residuals.

#### Importance of individual LIDAR-based measures and their relation to locally measured environmental factors

To evaluate the importance of the individual LIDAR measures we constructed an importance measure based on Akaike weights following Johnson and Omland (2004). To account for the fact that some variables were only allowed into a model once (if highly correlated with other predictors), and others were evaluated in many or all models, we had to modify the importance measure for each predictor. Therefore, initially each standardized coefficient was weighted with the model’s Akaike weight following Johnson and Omland (2004), summed and then finally this resulting value was weighted by a *predictor weight*. This predictor weight was the number of times the variable was retained in a model after AIC selection divided by the number of times the variable was allowed into a model. The absolute value of this weighted sum reflects the overall importance of each of the predictors for each species group, and will be referred to as the absolute importance in the following.

To evaluate the degree to which each of the 32 predictors can be used as proxies for any of the 15 measured environmental factors, we conducted pairwise Spearman’s rank correlations between these two sets of variables.

All statistical analyses were conducted in R version 3.4.2 (R Core Team, 2017).

## RESULTS

### LIDAR-biodiversity relations

We found that LIDAR-based measures have considerable predictive power for species richness in all species groups investigated (Table 2). Our best models, which had 4–7 LIDAR measures as predictors, yielded explanatory powers (R^2^) of 49, 31, 32 and 28 % for fungi, lichens, bryophytes and plants respectively. For all model details, see Table S4.4–S4.7.

**Table 2.** Best-model (based on highest cross validation score) and relative variable importance details. If a standardized coefficient (coef.) is given, the predictor in question was included in the best model for that particular species group. Since exclusion from the best models does not imply that a predictor is not important for the diversity of a specific species group, also absolute importance values (abs. imp.) of the most important predictors in the study (having abs. imp. ≥ 0.02 for at least one species group) are shown. The abs. imp. values of the three most important predictors are marked with bold. All predictors that are among these three most important for at least one species group are also highlighted in bold. All details on the modelling results are available in Table S2 in the supplementary material.

|  |  | **Macrofungi** |  | **Lichens** |  | **Bryophytes** |  | **Plants** |  |
| --- | --- | --- | --- | --- | --- | --- | --- | --- | --- |
|  | Meas. # | Coef. | Abs. imp. | Coef. | Abs. imp. | Coef. | Abs. imp. | Coef. | Abs. imp. |
| **Vegetation structures** |  |  |  |  |  |  |  |  |  |
| Point amplitude | 1 |  | 0.037 | -0.29 | 0.219 | -0.14 | 0.129 | 0.10 | 0.073 |
| **Point amplitude (ENT)** | 1 |  | 0.017 | 0.18 | 0.120 | 0.21 | **0.209** |  | 0.000 |
| **Biomass (ENT)** | 14 |  | 0.000 |  | 0.000 |  | 0.000 | -0.13 | **0.080** |
| Canopy openness (VAR) | 18 |  | 0.000 | 0.25 | 0.224 |  | 0.003 |  | 0.001 |
| Crown span | 16 | 0.26 | 0.003 |  | 0.000 |  | 0.000 |  | 0.000 |
| **Layer density (1.5 – 5 m)** | 7 | 0.31 | 0.011 | 0.35 | 0.016 |  | 0.002 | 0.28 | **0.257** |
| **Layer density (10 – 15 m)** | 9 |  | 0.000 |  | 0.000 |  | 0.001 | -0.22 | **0.222** |
| Layer density (25 – 30 m) | 12 |  | 0.011 |  | 0.000 | -0.09 | 0.026 |  | 0.000 |
| **Shrub layer height** | 17 |  | **0.380** |  | **0.356** | 0.24 | **0.218** |  | 0.021 |
| **Terrain structures** |  |  |  |  |  |  |  |  |  |
| **Heat load index (VAR)** | 21 | 0.14 | **0.125** | 0.41 | **0.411** | 0.22 | **0.218** |  | 0.000 |
| Terrain roughness | 22 |  | 0.002 | -0.28 | 0.220 |  | 0.000 |  | 0.000 |
| **Terrain slope** | 19 | 0.22 | **0.317** | 0.50 | **0.597** | 0.14 | 0.130 |  | 0.001 |
| Model # |  | 13 |  | 16 |  | 18 |  | 16 |  |
| Cross validation score |  | 0.81 |  | 0.59 |  | 0.54 |  | 0.38 |  |
| R<sup>2</sup> |  | 0.49 |  | 0.31 |  | 0.32 |  | 0.28 |  |

### The relative importance of LIDAR measures for biodiversity

Three LIDAR measures were important for three of the four species groups (fungi, lichens, and bryophytes): variation in local heat load, terrain slope and shrub layer height (Table 2). These were all positively related to local diversity. In addition to these three, four other LIDAR measures (i.e. seven in total) were ranked among the three most important for at least one of the species groups: point amplitude entropy, shrub layer density (1.5 – 5 m), medium-tree layer density (10 – 15 m), and variation in biomass (Table 2). Generally, these measures were also included in the best models for each species group (Table 2). While some of the measures were important for multiple species groups, some showed importance for only one or two species groups. These are detailed in Table 2 and illustrated in Fig. 3.

**Figure 3.**
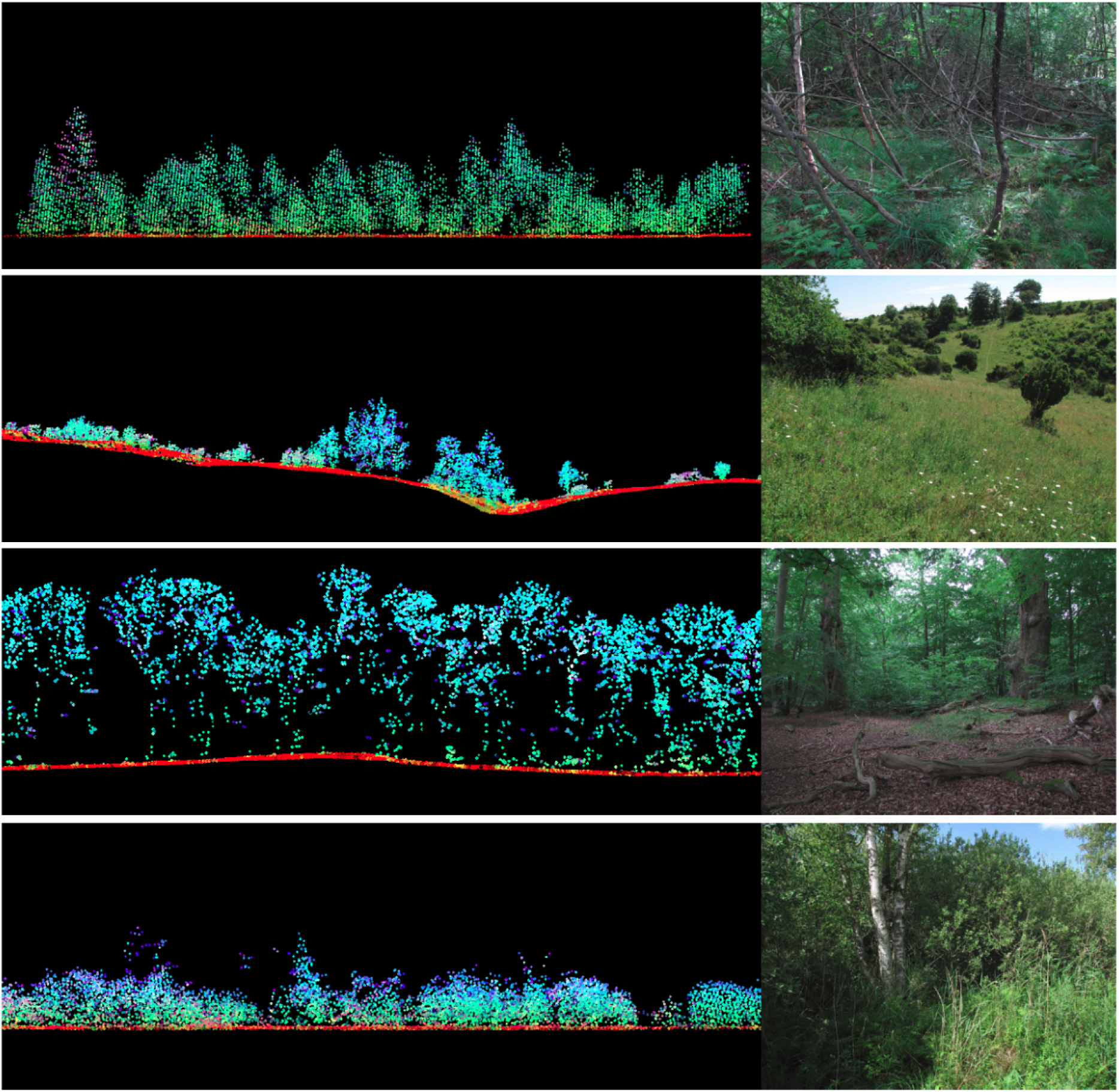
LIDAR point cloud cross sections and field photographs of characteristic species-rich locations for the four species groups. High species richness for bryophytes (top panel) was related to relatively steep terrain with relatively wet soils and a dense shrub layer. Locations with high vascular plant species richness (second from top) were open areas with low variability in biomass and high density in the shrub (1.5 – 5m) layer and low density of trees. For macrofungi (third panel from top) species richness were highest in areas with steep terrain with relatively wet soils but variable soil moisture levels, and a high degree of typical features for old-growth forest such as large crown spans, dead wood, high litter mass, and dense understory. High species-richness sites for lichens (bottom panel) were found in steep areas with a tall understory and variable canopy openness and on relative dry soils with variable moisture levels but little micro-topographic variation.

The seven LIDAR measures ranked as most important for local biodiversity were strongly correlated to several of the locally measured environmental variables (Table 3). Generally, the most important LIDAR measures representing vegetation structure (i.e., vegetation density, understory height, and biomass variation) were strongest and negatively related to the measured diurnal temperature differences and local light conditions. These LIDAR measures were also strongest and positively related to measurements of biotic factors such as litter mass, the volume of dead wood, and the coverage of old trees in the study sites (Table 3). The terrain LIDAR measures (i.e., local heat balance and terrain slope) were mainly related to locally measured soil moisture (Table 3). Generally, these LIDAR measures showed weaker relationships to the measured environment than those reflecting vegetation structure (Table 3).

**Table 3.**
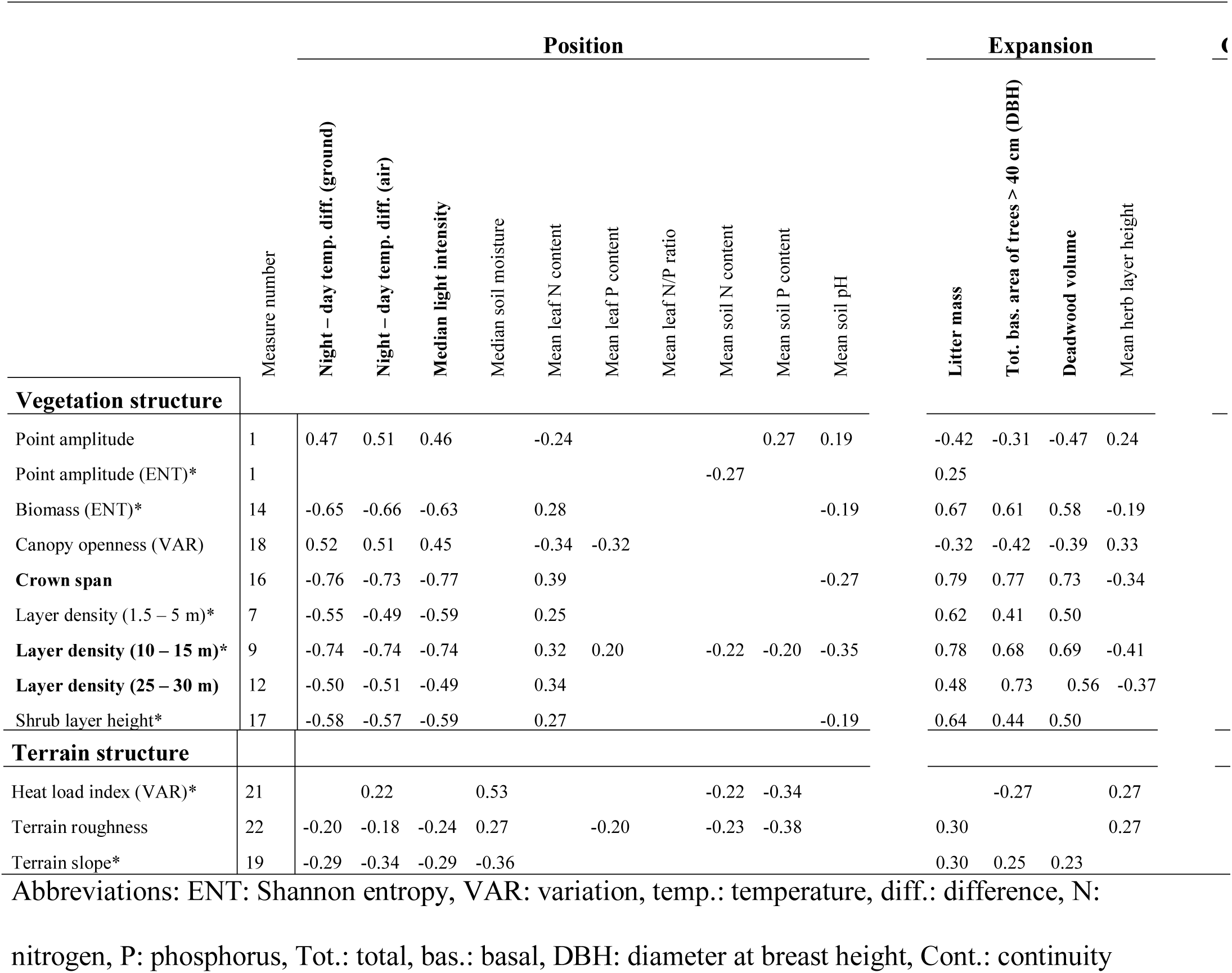
Spearman’s rho (only statistically significant values are shown, *p* < 0.05) of pairwise correlation analyses between the most important predictors in the study (having absolute importance (abs. imp.) ≥ 0.02 for at least one species group) and 15 environmental variables measured at each study site. The predictors and environmental variables in bold are those involved in at least one strong (rho > 0.7) relationship. Predictors marked with a “*” were ranked among the three most important for at least one of the species groups (see Table 1). The environmental variables are divided into the three components of ecospace (Brunbjerg et al. 2017b); position, expansion and continuity. The LIDAR measures are divided into those relating to vegetation and those reflecting terrain structures. The measure number relates to Table 1 where descriptions of the LIDAR measures can be found. No important (abs. imp. > 0.02) predictors were significantly correlated with local estimates of continuity; therefore, this column is empty.

The LIDAR measures most strongly correlated to local measurements of both abiotic and biotic factors (Spearman’s rho > 0.7) were the span of tree crowns and the vegetation density in the medium-tree (10 – 15 m) to upper (up to 30 m) height layers (Table 3). We also note that some of the factors typically thought to be important for local biodiversity such as soil moisture and vegetation height, were actually quite strongly related to a couple of our LIDAR measures (e.g., the topographic wetness index, Table S2). However, these were not among the most important LIDAR measures for local biodiversity identified in this study.

## DISCUSSION

To our knowledge, this is the first study demonstrating the suitability of LIDAR-based measures for predicting local (i.e., a few decameters) biodiversity patterns across several species groups and across all major temperate terrestrial ecosystems including fields, grasslands, wetlands, heathlands, dunes, scrubs and forests. So far, only a few studies have studied the extent to which LIDAR measures can predict diversity across multiple species groups, and these have included only one habitat type (see for example Zellweger et al., 2015 and Zellweger et al., 2016). We found that LIDAR measures alone can explain a considerable amount of local variation in biodiversity. A similar set of LIDAR measures were important for both bryophyte and lichen diversity but for fungi and plant diversity the set of important LIDAR measures differed notably. Hence, sites with high local species richness were structurally different in several aspects depending on the species group in question. Underpinning the importance of LIDAR for ecological monitoring, we found that the most important LIDAR measures were strongly related to both locally measured terrain and vegetation structures and can contribute to quantify expansion and partly position, while none of the important measures were related to continuity.

### The ability of LIDAR to explain local species richness

LIDAR measures could explain the diversity of macrofungi considerably better than the diversity of the other species groups. Although grasslands can hold quite a number of fungi species (Heilmann-Clausen and Vesterholt 2008), this group of organisms is notoriously known for its strong associations to old-growth structures, with old forests typically holding many species of macrofungi (Heilmann-Clausen and Vesterholt 2008). These forest structures are well represented by LIDAR derived measures and also known to be important for the diversity of plants, lichens and bryophytes (Camathias, Bergamini, Küchler, Stofer, & Baltensweiler, 2013; Lopatin, Dolos, Hernández, Galleguillos, & Fassnacht, 2016; Mao et al., 2018; Zellweger et al., 2015). However, for these groups the importance of terrain structures, microclimate, and soil-related factors are generally found to be more important than vegetation structures (Camathias et al., 2013; Zellweger et al., 2015). In particular, the local diversity in these groups strongly depend on soil characteristics (Ejrnæs and Bruun 2000, Ilomets et al. 2010, Ódor et al. 2013) and these are not well represented by LIDAR. This could explain the differences in predictive power between the species groups we observed here.

Intuitively, one could expect LIDAR to predict local diversity in forests better than in open landscapes since LIDAR represents the more complex 3D vegetation structure (biotic expansion) in forests particularly well. However, in our study the explanatory power obtained for all species groups corresponds well to – or is even better than – results from earlier studies relating LIDAR measures to species richness of fungi, lichens, plants and bryophytes (e.g., Camathias et al., 2013, Moeslund et al., 2013a, Thers et al., 2017, Bartels et al., 2018). This suggests that LIDAR is not only suitable for management and planning of diversity in forests, but is probably more broadly applicable and likely to be a valuable support tool for nature management and planning in open landscapes as well.

### Importance of individual LIDAR measures and their relation to locally measured environmental factors

Overall, the most important LIDAR measures for local biodiversity represented both vegetation (shrub layer height, point amplitude entropy, variation in biomass, shrub and medium-tree layer density) and terrain structures (slope of the terrain and variation in local heat load). The two terrain-structure measures correlated mostly with local soil moisture conditions, while the vegetation-structure measures were mainly associated with local light conditions and diurnal temperature variations, as well as biotic factors such as litter mass, stand age and the amount of dead wood.

In previous studies, local terrain structure have been shown to affect both the occurrence, abundance and species richness of macrofungi (Peura et al. 2016, Thers et al. 2017, Chen et al. 2018). Here, we found fungal species richness to be highest in areas with steep terrain with relatively wet soils, while at the same time having many typical features for old-growth forest such as large crown spans, large amounts of dead wood, high litter mass, and a dense shrub layer. This supports that this group is typically strongly associated with old-growth structures (Heilmann-Clausen and Vesterholt 2008) and suggests that steep slopes in this case could reflect refugia from human impact (Odgaard et al. 2014).

The most important LIDAR measures for local diversity of bryophytes and lichens were almost similar. This hints that in nature the same natural factors determine the local diversity patterns of these two species groups – a finding that was also highlighted by Pharo and Beattie (1997). However, for lichens terrain slope was the most important predictor and had a strong positive relationship to local species richness, whereas for bryophytes, variation in heat load index was an important predictor. Since, we found terrain slope negatively, and variation in heat load index positively related to locally measured soil moisture, we believe these results support (Pharo and Beattie 1997); bryophyte richness is higher in relatively moist sites and lichen richness is higher in the drier sites. For bryophytes and lichens, local diversity decreased with mean point amplitude and increased with point amplitude variation. As point amplitude can be interpreted as a measure of successional stage from bare soil (high point amplitude) to closed forest (low point amplitude, see methods), this indicates that species richness of bryophytes and lichens is often higher in late successional stages (old growth forests and old scrubland). Furthermore, local diversity of both groups was positively related to shrub layer height, which were associated with light availability, microclimatic conditions, litter mass, stand age and the amount of dead wood. These findings correspond to the current knowledge based on single-habitat studies. For example, Mills and Macdonald (2004) and Zellweger et al. (2015) showed that microsite bryophyte diversity in forests was clearly affected by dead wood characteristics, and local levels of soil moisture, temperature and solar radiation among others. Similarly, Leppik et al. (2013) found that forest lichen diversity increased with stand age and soil moisture. Note that while we assess the importance of LIDAR measures for all lichens and bryophytes, considerable differences in predictor importance can be expected among edaphic and epiphytic species (e.g., Camathias et al., 2013).

For vascular plants, we found high species richness at localities with high density in the shrub layer and low density of medium sized trees, and in areas with low variability in biomass and a high mean point amplitude (indicating early successional stage, i.e. open landscapes). These results suggest that plant diversity is often high in open landscapes, e.g. grasslands, which are known hotspots for plants in Northern Europe (Habel et al., 2013). On the other hand, they also indicate that areas with relatively many shrubs or small tress are rich in plant species. A combination of two processes might explain this. First of all, grasslands are threatened by shrub encroachment (Timmermann et al. 2015) and the diversity-density relationship could therefore reflect an extinction debt to unfavorable habitat conditions following encroachment. Secondly, shrubs create additional microhabitats in open grasslands and could thereby increase richness. The later supports that plant diversity can be promoted by the presence of single-standing trees and bushes in otherwise homogenous grassland swards (Moeslund et al. 2017). Contrary to previous findings, we did not find evidence that terrain-related factors are important for determining the species richness of plants. For example, terrain controlled soil moisture has been found important for local diversity of plants in open habitats and is generally regarded as important for plant species richness (Moeslund et al. 2013a, Silvertown et al. 2015). However, we found no clear indications of a relationship between soil moisture and plant diversity. We included forests in this study and here soil moisture is probably less important for shaping local vegetation patterns (Zellweger et al. 2015) due to the more moist local climate mediated by trees. This may explain the lack of this otherwise important relationship in the present study. However, local environmental variation unaccounted for by LiDAR might also mask the effect of soil moisture. Exploring this in more detail could be the focus of future studies.

Our LIDAR measures captured major aspects of the environmental variation related to build up and diversification of organic matter (*ecospace expansion* sensu Brunbjerg et al., 2017b). For example, the measured litter mass, the area with old trees (stem diameter at breast height > 40 cm) and the volume of dead wood were all highly correlated with at least one of our LIDAR measures. Several studies have reached a similar conclusion in forests (Camathias et al. 2013, Zellweger et al. 2015, Lopatin et al. 2016). However, our results demonstrate for the first time that LIDAR can be used to estimate expansion-related factors along the full successional gradient from open wetlands, grasslands and fields to scrubs and forests. This opens interesting perspectives for applying LIDAR more broadly in nature management and planning (see section on Application perspectives).

The LIDAR measures used in this study were good representatives of soil moisture and local temperature conditions, but failed to act as proxies for other ecospace position related factors such as soil pH and nutrient status. By nature, LIDAR does not record anything below ground, nor any chemical properties. This renders LIDAR alone unsuitable for recording soil and leaf chemistry, but there could be differences in terrain and vegetation structure across habitat types mirroring these factors of which we are unaware. Denser point clouds combined with new machine learning techniques (Liu et al. 2018) and perhaps spectral information (Lausch et al. 2016) could potentially remedy this situation in the future.

Our LIDAR measures were generally quite poor proxies for ecospace continuity (Brunbjerg et al. 2017b). Since spatial continuity by definition is a broad-scale factor (Nordén et al. 2014), it is not surprising that this factor was poorly represented by our LIDAR measures. Though, by considering LIDAR point clouds for a larger area than we did in this study, we believe LIDAR could be valuable for estimating spatial continuity and we urge researchers to attempt to do this in future studies. On the other hand, temporal continuity is tricky to estimate from LIDAR, at least in open landscapes where the structures and vegetation patterns characteristic of long continuity are difficult to capture. For example considerable plant species turnover can occur over time from abandoned fields to heathland and grassland, without noticeable changes in vegetation structure (Ejrnæs et al. 2008). Contrary, in forests – as we have shown – some of the structures characteristic of old-growth can be estimated with high confidence using LIDAR. Clearly, more work and possibly technological advances are needed to find methods to effectively estimate temporal continuity using LIDAR.

### Application perspectives

Using LIDAR, researchers and managers have gained the ability to cover large areas (even whole nations) in adequately fine detail for nature planning and management (McElhinny et al. 2005). In fact, LIDAR has the potential to play a crucial role in this applied field by enabling detailed mapping and assessments for decision makers and field biologists, while at the same time covering the desired extent. The best candidates for LIDAR measures with potential for supporting conservation planning and management are those being notoriously difficult to quantify in the field, having high importance across species groups and a plausible ecological interpretation. Generally, a measure such as shrub layer height is indeed complicated and time-consuming to quantify in the field, but had high importance for most species groups in this study and seemed to capture several of the factors of importance for local biodiversity patterns (see discussion above). Another such measure is the terrain aspect-based heat load index, which proved to represent soil moisture well and be important for most species groups in our study. While more accurate terrain-based wetness measures exist (Hengl and Reuter 2009, Moeslund et al. 2013a), this indicator has the advantage that it is computationally efficient since it can be calculated without the need to delineate watersheds. Hence, these LIDAR measures are two out of potentially several that may be successfully implemented in planning and management, and used to – for example – create quick first-impressions of local biodiversity patterns across large areas in the future.

In summary, our results show that LIDAR alone can provide reasonable predictive power for biodiversity, giving insights into local biodiversity patterns and their potential drivers. By refining these methods, for example by (1) including full-waveform LIDAR (Anderson et al. 2016), (2) further investigate LIDAR-based measures for assessing continuity and maybe soil chemistry (see discussion above) and (3) possibly combining LIDAR with other available data and methods, this technology could open new avenues offering rigid fine-grain overviews of the biodiversity patterns and potentially also dynamics for whole countries. This would allow hitherto unseen possibilities for evidence-based biodiversity management (Brunbjerg et al. 2016, Mao et al. 2018), and help to precisely target field-based biodiversity monitoring nationwide.

## ACKNOWLEDGMENTS

We sincerely thank Villum Fonden for funding the Biowide project. We gratefully acknowledge the contributions from those who collected field data for Biowide: macrofungi (Thomas Læssøe), lichens (Ulrik Søchting and Roar Poulsen), bryophytes (Irina Goldberg) and plants (Irina Goldberg, Peter Wind, Hans Henrik Bruun). JEM was supported by Aage V. Jensen Naturfond. SN and AZ were supported by Aarhus University Research Foundation (AUFF-E-2015-FLS-8-73), JCS and AZ by the European Research Council (ERC-2012-StG-310886-HISTFUNC), and AZ by the Hungarian Research Fund (OTKA PD 115833 and ERC_16_M 122670). JEM considers this work a contribution to the project “Dark Diversity in nature management” funded by Aage V. Jensen Naturfond, SN considers it a contribution to her Carlsberg Distinguished Associated Professor Fellowship “Sensing biodiversity change and its drivers”, and JCS considers it a contribution to his Villum Investigator project “Biodiversity Dynamics in a Changing World” funded by Villum Fonden.

