## Appendix 1 for "LIDAR explains diversity of plants, fungi, lichens and bryophytes across multiple habitats and large geographic extent"

**Appendix 1.** The full OPALS script used for the study.

::Amplitude mean, entropy, variance, RMS

opalscell -infile *.odm -attribute amplitude -feature mean entropy rms variance -cellsize 10

::Echo number RMS

opalscell -infile *.odm -cellsize 10 -feature mean rms -attribute EchoNumber

::Terrain slope and aspect

opalsgridfeature -infile DTM_*tif -feature slope exposition -kernelsize 1
opalsstatfilter -inf *slope.tif -feature mean -kernelsize 12 -kernelshape square -gridsize kernelsize

::DTM heat load index, DTM heat load index variance, entropy

opalsalgebra -infile *expos.tif -formula "(1-cos(r[0]-0.785398))/2")
opalsstatfilter -inf *algebra.tif -feature mean -kernelsize 12 -kernelshape square -gridsize kernelsize)

opalsstatfilter -inf *algebra.tif -feature variance entropy -kernelsize 12 -kernelshape square -gridsize kernelsize)

::Roughness of surface (Sigma Z)

opalsgrid -infile *.odm -filter "Class[ground]" -interp movingplanes -gridsize 0.5 -searchrad 0.75 -neighbours 6 -feature sigmaz

::Local terrain openness

opalsopenness -inf DTM_hi_res*.tif -outf *_avg_openness.tif -feature positive -kernelsize 10 -selmode 0
opalsstatfilter -inf *_avg_openness.tif -outf *_DTM_openness_mean.tif -feature mean -kernelshape square -kernelsize 10 -gridsize kernelsize

::Landscape-scale terrain openness

opalsopenness -inf *_10m_dtm.tif -feature positive -selmode 0 -kernelsize 15 -outfile *DTM_landscape_openness.tif

::dtm_openness_diffminmax_mean
opalsopenness -inf DTM_hi_res_*.tif -outf *_min_openness.tif -feature positive -kernelsize 10 -selmode 1
opalsopenness -inf DTM_hi_res_*.tif -outf *_max_openness.tif -feature positive -kernelsize 10 -selmode 2

opalsalgebra -inf *_min_openness.tif *_max_openness.tif -outf *_diff_openness.tif -formula "r[0]-r[1]"

opalsstatfilter -inf *_diff_openness.tif -outf *_DTM_openness_diffminmax_mean.tif -feature mean -kernelshape square -kernelsize 10 -gridsize kernelsize

::TWI: from already existing dataset

::normalized height

opalsaddinfo -gridfile DTM_hi_res_*.tif -infile *.odm -attribute "NormalizedZ = z - r[0]"

opalscell -infile *.odm -attribute NormalizedZ -feature mean -cellsize 10

opalscell -infile *.odm -attribute NormalizedZ -feature range -cellsize 10

opalscell -infile *.odm -attribute NormalizedZ -feature entropy -cellsize 10

opalscell -infile *.odm -attribute NormalizedZ -feature rms -cellsize 10

::Echo ratio (excluding building points)

opalsnormals -infile *.odm -normalsalg fmcd -neighbours 8 -searchrad 1.5 -storemetainfo medium)
opalsechoratio -infile *.odm -filter "Class[LowVegetation MediumVegetation HighVegetation Ground unclassified]" -searchrad 1.5 -ratiomode slopeAdaptive

opalscell -infile *.odm -attribute EchoRatio -feature mean entropy variance -cellsize 10

::point count in height classes

::pcount_1.5_5
opalscell -inf *.odm -outf *_pc_1.tif -feat pcount -cells 10 -filter "generic[normalizedz >= 1.5 and normalizedZ < 5]"

::pcount_5_10
opalscell -inf *.odm -outf *_pc_2.tif -feat pcount -cells 10 -filter "generic[normalizedz >= 5 and normalizedZ < 10]"

::pcount_10_15
opalscell -inf *.odm -outf *_pc_3.tif -feat pcount -cells 10 -filter "generic[normalizedz >= 10 and normalizedZ < 15]"

::pcount_15_20
opalscell -inf *.odm -outf *_pc_4.tif -feat pcount -cells 10 -filter "generic[normalizedz >= 15 and normalizedZ < 20]"

::pcount_20_25
opalscell -inf *.odm -outf *_pc_5.tif -feat pcount -cells 10 -filter "generic[normalizedz >= 20 and normalizedZ < 25]"

pcount_25_30
opalscell -inf *.odm -outf *_pc_6.tif -feat pcount -cells 10 -filter "generic[normalizedz >= 25 and normalizedZ < 30]"

::pcount_30_inf
opalscell -inf *.odm -outf *_pc_7.tif -feat pcount -cells 10 -filter "generic[normalizedz >= 30 ]"

:: calculate the sum of all points for layer count

opalsalgebra -inf %1_pc_1.tif %1_pc_2.tif %1_pc_3.tif %1_pc_4.tif %1_pc_5.tif %1_pc_6.tif %1_pc_7.tif -outf %1_pc_0.tif -formula max(r) -resamp nearest

:: normalize the point count and binarize the data for layer count

set thr=0.03

opalsalgebra -inf %1_pc_1.tif %1_pc_0.tif -outf %1_pc_1_bin.tif -formula "(r[0]/r[1]) > %thr%*1 ? 1 : 0" -resampling nearest

opalsalgebra -inf %1_pc_2.tif %1_pc_0.tif -outf %1_pc_2_bin.tif -formula "(r[0]/r[1]) > %thr%*1 ? 1 : 0" -resampling nearest

opalsalgebra -inf %1_pc_3.tif %1_pc_0.tif -outf %1_pc_3_bin.tif -formula "(r[0]/r[1]) > %thr%*1 ? 1 : 0" -resampling nearest

opalsalgebra -inf %1_pc_4.tif %1_pc_0.tif -outf %1_pc_4_bin.tif -formula "(r[0]/r[1]) > %thr%*1 ? 1 : 0" -resampling nearest

opalsalgebra -inf %1_pc_5.tif %1_pc_0.tif -outf %1_pc_5_bin.tif -formula "(r[0]/r[1]) > %thr%*2 ? 1 : 0" -resampling nearest

opalsalgebra -inf %1_pc_6.tif %1_pc_0.tif -outf %1_pc_6_bin.tif -formula "(r[0]/r[1]) > %thr%*2 ? 1 : 0" -resampling nearest

opalsalgebra -inf %1_pc_7.tif %1_pc_0.tif -outf %1_pc_7_bin.tif -formula "(r[0]/r[1]) > %thr%*2 ? 1 : 0" -resampling nearest

:: detect changes between neigbouring layers for layer count

algebra -inf %1_pc_1_bin.tif	-outf %1_L1_con.tif -formula "r[0] > 0 ? 1 : 0"	-resampling nearest

algebra -inf %1_pc_1_bin.tif %1_pc_2_bin.tif -outf %1_L2_con.tif -formula "r[0]<r[1] ? 1 : 0" -resampling nearest

algebra -inf %1_pc_2_bin.tif %1_pc_3_bin.tif -outf %1_L3_con.tif -formula "r[0]<r[1] ? 1 : 0" -resampling nearest

algebra -inf %1_pc_3_bin.tif %1_pc_4_bin.tif -outf %1_L4_con.tif -formula "r[0]<r[1] ? 1 : 0" -resampling nearest

algebra -inf %1_pc_4_bin.tif %1_pc_5_bin.tif -outf %1_L5_con.tif -formula "r[0]<r[1] ? 1 : 0" -resampling nearest

algebra -inf %1_pc_5_bin.tif %1_pc_6_bin.tif -outf %1_L6_con.tif -formula "r[0]<r[1] ? 1 : 0" -resampling nearest

algebra -inf %1_pc_6_bin.tif %1_pc_7_bin.tif -outf %1_L7_con.tif -formula "r[0]<r[1] ? 1 : 0" -resampling nearest

:: derive the number of layers

algebra -inf %1_L1_con.tif %1_L2_con.tif %1_L3_con.tif %1_L4_con.tif %1_L5_con.tif %1_L6_con.tif %1_L7_con.tif -outf %1_layer_count.tif -formula sum(r) -resampling nearest

::pseudowaveform, pseudowaveform variance

opalspointstats -infile *.odm -filter "Class[Ground Lowvegetation mediumvegetation highvegetation unclassified]" -searchmode d2_5 -searchrad 0.5 1 -feature variance -refmodel tiltedplane
opalscell -infile *.odm -feature variance entropy -cellsize 10 -attribute _dZVariance

::d_biomass, d_biomass_entropy

opalsaddinfo -infile *.odm -attribute "_d_biomass = (Abs(NormalizedZ) / 3) + (Abs(NormalizedZ) * Echoratio / 36) + (Abs(NormalizedZ) * NrofEchos / 6)" -filter "Class[unclassified ground lowvegetation mediumvegetation highvegetation]"

opalscell -infile *.odm -attribute NormalizedZ -feature entropy -cellsize 10

::canopy top height mean, canopy base height mean

opalscell -inf %1.odm -cells 2.5 -attri normalizedz -feature quantile:0.90 quantile:0.05 -filter "generic[normalizedz >= 3 and normalizedz < 50]"

opalsstatfilter -inf %*_NormalizedZ_q0.90.tif -outf *_canopy_top_10m.tif -feature mean -kernelsize 2 -gridsize kernelsize -kernelshape square

opalsstatfilter -inf %*_NormalizedZ_q0.05.tif -outf *_canopy_base_10m.tif -feature mean -kernelsize 2 -gridsize kernelsize -kernelshape square

::understorey_height_mean
opalscell -inf *.odm -cellsize 2.5 -attribute NormalizedZ -feature quantile:0.90 -filter "generic[normalizedz > 0.3 and normalizedz < 3]"
opalsstatfilter -inf *_NormalizedZ_q0.90.tif -outf *_ust_height_10m.tif -feature mean -kernelsize 2 -gridsize kernelsize -kernelshape square

::crown span

algebra -inf %1_NormalizedZ_q0.90.tif %1_NormalizedZ_q0.05.tif -outf %1_crown_height_2_5m.tif -formula "r[0]-r[1]"

opalsstatfilter -inf %1_crown_height_2_5m.tif -outf %1_crown_span_mean.tif -feature mean -kernelsize 2 -gridsize kernelsize -kernelshape square

::canopy openness mean and variance

canopy_openness_mean
opalspointstats -infile *.odm -feature posopenness -searchrad 5 -filter "Class[unclassified ground lowvegetation mediumvegetation highvegetation
opalscell -inf *.odm -attribute _dZPosOpenness -feature median -cellsize 0.5 -filter "Echo[Last]"
opalsstatfilter -inf *_dZPosOpenness_median.tif -feature mean -kernelsize 10 -kernelshape square –gridsize kernelsize

opalsstatfilter -inf *_dZPosOpenness_median.tif -feature variance entropy -kernelsize 10 -kernelshape square –gridsize kernelsize

::crown span mean
opalsalgebra -inf *_NormalizedZ_q0.90.tif *_NormalizedZ_q0.05.tif -outf *_crown_height_2_5m.tif -formula "r[0]-r[1]"
opalsstatfilter -inf %1_crown_height_2_5m.tif -outf *_crown_height_10m.tif -feature mean -kernelsize 2 -gridsize kernelsize -kernelshape square
