## Supplementary material for "LIDAR explains diversity of plants, fungi, lichens and bryophytes across multiple habitats and large geographic extent": Fig. S3

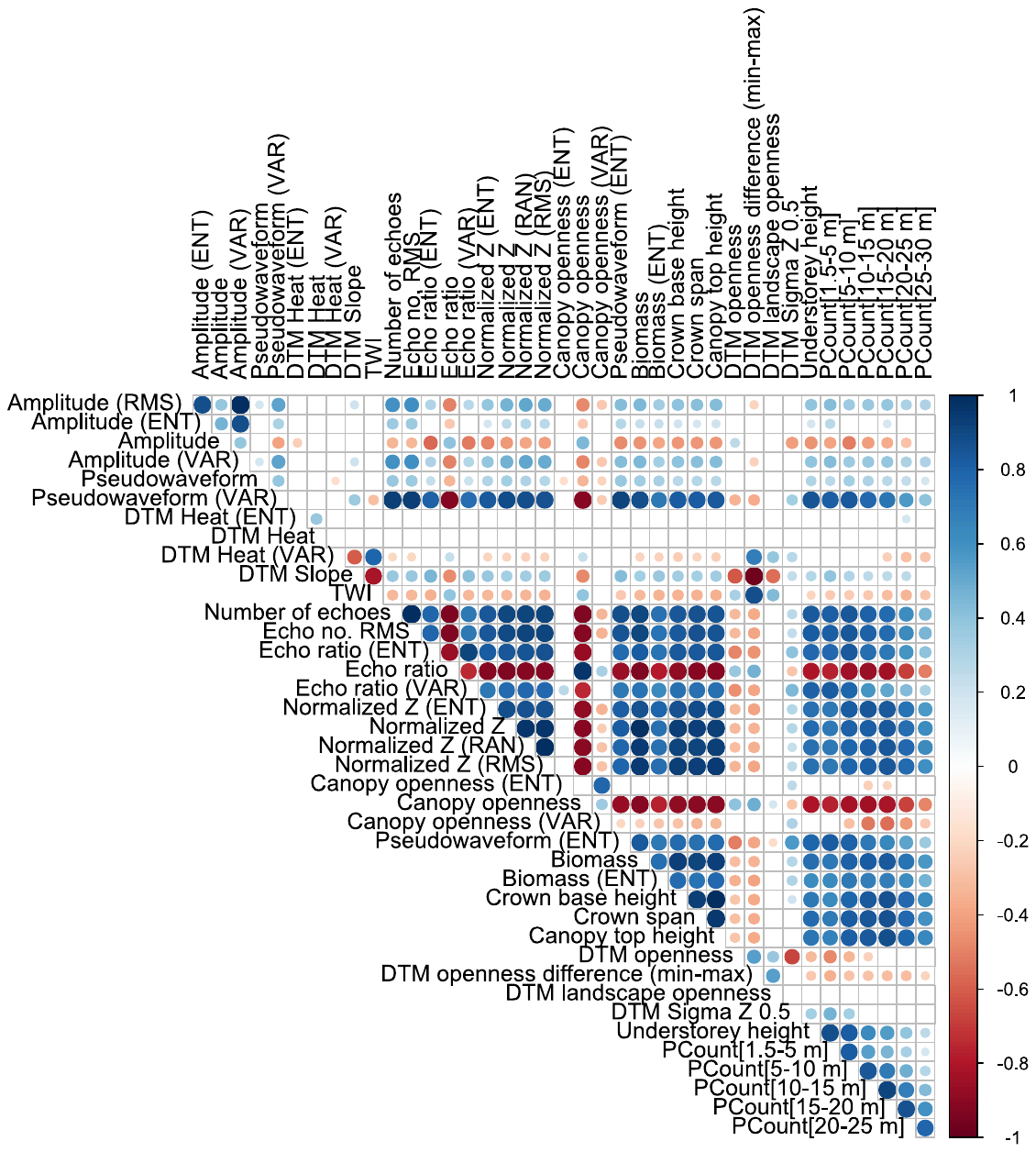

Fig. S3. Pairwise Spearman’s correlation coefficients (rho) between all LIDAR measures and their calculated variance measures. Only statistically significant correlations are shown (*p* < 0.05). Abbreviations: ENT: Shannon entropy, RAN: Range, VAR: Variance, RMS: Root mean square error, DTM: digital terrain model
